# Predator sounds disrupt visual attention in jumping spiders

**DOI:** 10.64898/2026.10.21.760757

**Authors:** Alex M. Winsor, Guilherme F. Pagoti, Daniel Daye, Philip Denbaum, Ronald R. Hoy, Elizabeth M. Jakob

**Affiliations:** Organismic and Evolutionary Biology, University of Massachusetts Amherst, Amherst, MA 01003, USA; Programa de Pós-Graduação em Zoologia, Instituto de Biociências, Universidade de São Paulo, Rua do Matão, 321, Travessa 14, Cidade Universitária, São Paulo - SP, 05508-090 Brasil; Department of Biology, University of Massachusetts Amherst, Amherst, MA, 01003, USA; Department of Neurobiology and Behavior, Cornell University, Ithaca, NY 14853, USA

## Abstract

Perception in one sensory modality can influence not only what animals attend to in another, but also whether animals increase their vigilance or remain engaged with ongoing sensory stimuli at all. Jumping spiders (Salticidae) are highly visual animals, but they are also alert to auditory cues. Previous work has shown that jumping spiders respond to the sound of the wingbeats of wasps, a potential predator, by freezing. Here, we examined whether wasp sounds influence eye movements, an indicator of visual attention. We used a custom-built eye tracker to measure the gaze direction of the movable, high-acuity principal eyes, and tested whether spiders would shift their gaze toward the location of a threat. In Experiment 1, when spiders were played the sound of a buzzing wasp, their retinas, initially at rest, immediately began sweeping rapidly back and forth, but were not directed toward the location of the sound. In Experiment 2, spiders scanning a salient prey stimulus (a cricket silhouette) with their principal eyes were simultaneously presented with a wasp sound and the appearance of a visual distractor. Compared to trials in which the same spiders were shown the distractor without sound, spiders were more likely to shift their gaze away from the cricket but were no more likely to redirect their gaze toward the distractor. Because acoustic cues from aerial predators may indicate immediate danger without providing reliable spatial information, orienting toward the wrong target may be costly under high uncertainty. Our results demonstrate that predator sounds do not spatially guide visual attention in jumping spiders, but instead disrupt attention to salient visual stimuli by increasing non-directed eye movements.

## INTRODUCTION

Information from one sensory modality can influence how animals interpret stimuli in another sensory modality. This cross-modal interaction helps determine whether animals stay engaged in an ongoing behavior, or instead shift to another behavior or a state of heightened alertness. When signals or cues across modalities are predictable and mutually reinforcing, they can help support decision-making and sustain attention (Partan and Marler, 1999; Hebets and Papaj, 2013). In contrast, those that are highly unexpected or salient can interrupt ongoing processing and redirect attention toward higher-priority information. Animals must often integrate information across their senses to reduce uncertainty. For example, combined multisensory cues can improve accuracy in predator recognition (Ward and Mehner, 2010) and social decisions (Munoz and Blumstein, 2020).

Predator-associated signals, such as alarm calls produced by social partners, and predator-associated cues, including odors or sounds generated by the predator itself, can disrupt ongoing attention to promote broad scanning of the environment. This reflects a shift toward increased vigilance (Lima and Dill, 1990; Lima, 1998). Such shifts often come at the cost of suspended engagement with ongoing behaviors like foraging or hunting, yet they can enhance survival by enabling earlier detection of threats and faster initiation of escape responses (Blumstein, 2006). In these cases, inputs from more than one sensory modality may increase the likelihood that attention is redirected away from ongoing tasks and toward potential threats. In other cases, sudden stimuli can instead elicit a startle response that briefly interrupts ongoing behavior, rather than redirecting attention toward any particular stimulus (Yeomans and Frankland, 1995).

Here we examine how auditory cues influence visual perception in jumping spiders. Jumping spiders rely on specialized eyes that support remarkable vision despite their small size, helping them to assess elaborate courtship displays (e.g., Taylor and McGraw, 2013; Painting et al., 2016), as well as the color patterns of potential prey (Taylor et al., 2014) and predators (Rößler et al., 2021). Their visual system is distributed across eight eyes belonging to two functional classes. Three pairs of secondary eyes positioned around the cephalothorax act primarily as wide-field motion detectors. One of these pairs, the anterior lateral eyes (ALEs), faces forward and shares overlapping visual fields with the principal eyes. The principal eyes possess narrow, boomerang-shaped retinas densely packed with photoreceptors (reviewed in Harland et al., 2012; Morehouse, 2020; Winsor et al., 2024). Detection of a stimulus by the ALEs triggers the spider to direct its principal eyes toward the target (Jakob et al., 2018), analogous to how some vertebrates shift their gaze from peripheral vision to high-acuity foveal vision. Although the principal eyes have fixed external lenses and narrow fields of view, the principal-eye retinas reside within mobile tubes that allow spiders to internally move their eyes and inspect objects with high spatial resolution (Land, 1969a; 1969b).

Despite the prominence of vision, jumping spiders also possess a suite of well-developed nonvisual sensory modalities. In particular, they are highly sensitive to both substrate-borne and far-field airborne vibrations. For example, male spiders deliberately strum courtship songs into the substrate, which females use to assess male quality (e.g., Elias et al., 2003; 2005), and spiders can also exploit vibrations inadvertently produced by prey insects to capture them (Hergenröder and Barth, 1983). Although it has long been demonstrated that jumping spiders respond to vibrations, until recently it was less clear whether and how spiders could detect airborne pressure waves, as they lack tympanic membranes like those of other arthropods. Shamble et al. (2016) showed that jumping spiders in an anechoic chamber can detect airborne sound, specifically at low frequencies similar the buzz of a wasp, one of their key predators, and the authors recorded auditory evoked neural responses to this sound in a central brain region that supports sensorimotor integration for flexible behavior.

When jumping spiders hear wasp sounds, they freeze, presumably to avoid detection by the wasp (Shamble et al., 2016). We predicted that because the movements of the internally moving eyes are difficult for an observer to detect, a spider that detects a wasp sound should increase their eye movements and either look toward the sound source or upwards, the likely direction of wasp approach (Experiment 1). We also tested the influence of a wasp sound on visual attention. We predicted that upon hearing a wasp buzz, spiders inspecting an image of a cricket prey that normally captures their attention would look away from the prey and toward a suddenly appearing distractor (Experiment 2).

## METHODS

### Spider collection and housing

We conducted experiments using two closely related jumping spider species, *Phidippus princeps* and *P. audax*, which exhibit comparable life history traits, including mate-searching behavior, hunting strategies, and exposure to similar predators. Because availability of these species varied across seasons and years, the species used in each experiment depended on the timing of collection. Spiders were collected from multiple locations in western Massachusetts during late fall, with a cohort of *P. princeps* obtained in 2016 and a cohort of *P. audax* obtained in 2019. Individuals were captured using sweep nets in areas dominated by grasses and flowering plants (Hoefler and Jakob, 2006) and were collected in vials by hand from built structures such as fences and bridges. Most spiders were adults or penultimate instars at the time of capture, and all experimental subjects were field caught.

After collection, spiders were housed singly in plastic enclosures (18 x 13 × 10 cm) and maintained at approximately 25°C under a 16:8 h light: dark cycle with full-spectrum illumination. Spiders were fed banded crickets (*Gryllodes sigillatus*) once per week, with crickets reared on a high-protein dog food diet. Water was available continuously from cotton-plugged vials. To standardize environmental complexity, each enclosure contained a black plastic tube, artificial foliage, and a green-painted climbing dowel (Carducci and Jakob, 2000). All spider enclosures were kept on shelves in a relatively quiet laboratory.

### Eyetracking methods

In Experiments 1 and 2, we recorded gaze direction of the principal eyes using a custom-built eyetracking system designed for jumping spiders (Canavesi et al., 2011). During testing, spiders were tethered in front of the apparatus and viewed back-projected video stimuli. The principal-eye retinas were illuminated with infrared (IR) light, wavelengths outside the spiders’ visual sensitivity range, and the resulting retinal reflections were captured using an IR-sensitive camera. Following each experiment, retinal recordings were digitally aligned with the stimulus videos to quantify gaze direction with high precision (Figure 1).

**Figure 1.**
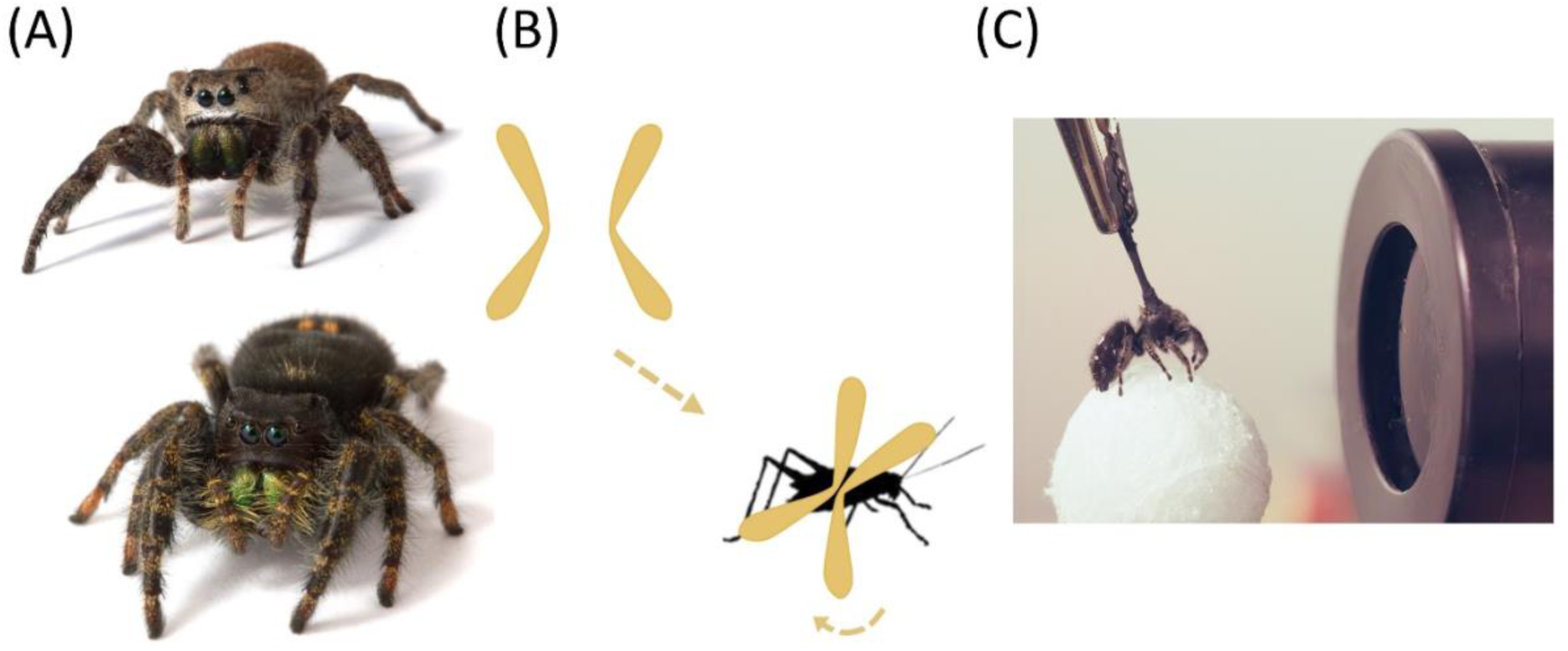
Experimental animals and behavioral assay designs. (A) Front-facing views of adult female *Phidippus princeps* (top) and *Phidippus audax* (bottom) jumping spiders, the species used in Experiments 1 and 2, respectively. The prominent central eye pair are the principal eyes, which provide high spatial resolution, and are bordered laterally by the anterior lateral eyes (ALEs), a pair of secondary eyes that are specialized for motion detection and overlap with the principal eyes in their fields of view. (B) The visual fields of the principal-eye retinas (yellow) are boomerang-shaped and coordinate their movements to examine objects of interest through shifting, scanning, and rotational motions. In this schematic, the retinas pivot to align with and scan a newly appearing cricket stimulus. (C) A tethered *P. audax* positioned within the eye-tracking apparatus, which enables precise measurement of gaze direction during stimulus presentation. Loudspeakers out of frame are positioned in an array around the spider.

IR illumination differed between experiments. In Experiment 1, we used an 850 nanometer (nm) IR light beam delivered via a dual-branch light guide fitted with a focusing lens (Thorlabs Inc., Newton, NJ, USA). In Experiment 2, we used a longer-wavelength 940 nm IR source with a focusing lens, which improved image quality. Retinal movements were recorded at 15 frames per second (fps) using an IR-sensitive IDS camera (Edmund Optics) controlled through the Image Processing Toolbox in MATLAB (version R2011b). Visual stimuli, when presented, were coded in the software Processing (version 3.5.4) and presented to the spider using a miniature projector (AAXA P4-X Pico), while a separate monitor displayed synchronized recordings of retinal position and stimulus presentation in real time. We used a screen recorder (OBS) which simultaneously recorded the IR retina camera feed and the visual stimulus projection at 30 fps.

Spiders were secured in the eyetracker following previously described tethering methods (Jakob et al., 2018; Bruce et al., 2021). Briefly, a 1:1 mixture of gum rosin (Acros Organics, Fairlawn, NJ, USA) and beeswax (Stackich Inc., Troy, MI, USA) was heated using a dental wax applicator, and the brush end of a microapplicator (Shintop disposable microbrushes, 1.5 mm) was dipped into the molten mixture. The microbrush was then attached to the spider’s cephalothorax (head), allowing reversible tethering. After the wax had rapidly hardened, the brush was coated with matte black paint to reduce stray reflections (Stuart Semple Black Paint, CultureHustle, Dorset, UK). The microbrush was clamped in position to orient the spider toward the eyetracker. To increase their comfort, spiders were provided with a lightweight polystyrene ball to grasp and “walk” on. Individuals were permitted to acclimate for 10 min prior to testing.

### Experiment 1: Retinal movement in response to predator sounds

We tested the response of field-caught adult female *P. princeps* spiders to an acoustic stimulus, the sound of a wasp, in the absence of visual cues. We confirmed in preliminary experiments that freely moving *P. princeps* froze in response to the sound in the same way as did the *P. audax* observed by Shamble et al. (2016). Three Bluetooth-enabled speakers (Sony SRS-X11) were placed on speaker stands (VideoSecu MS07B) in an array around the spider secured in the eyetracker. The speakers faced the spider. To ensure there was a clear line of sight between the spider and the speaker without being potentially blocked by the eyetracker and tethering apparatus, the speakers were positioned slightly higher and were angled downward at approximately 8° from vertical to face the spider. Speakers were each placed 53 cm away. One speaker was directly behind the spider and two front speakers were placed at angles of approximately 20° to the left and right of the spider, well within the approximately 50 ° field of view of the principal eyes (Land, 1969). The tethered spiders were thus able to direct the gaze of the principal eyes at the two front speakers, but not at the rear speaker. Each speaker was connected to an iPod Touch (Apple Inc., Cupertino, CA) via Bluetooth.

Once the spider was positioned in front of the eyetracker, the room was darkened, and the spiders (*N* = 13) were allowed to acclimate until spontaneous movement of the retinas stopped for at least 10 seconds. Once the retinas were still, we started the sound stimulus (Supplementary Video 1). We presented the sound from each of the three speakers in random order using a random number generator (random.org/lists). Each sound stimulus presentation lasted for 5 seconds and then followed by a pause of at least 30 seconds or until the retinas stopped moving for at least 10 seconds. For this experiment, spiders viewed a blank white screen because we were interested in retinal movements in the presence of sound but in the absence of visual stimuli.

Using Final Cut Pro, videos were downsampled to 10 fps and exported as a stack of JPEG images. Frame-by-frame tracking was performed using the ImageJ distribution FIJI (fiji.sc) and the MTrackJ plugin (https://imagescience.org/meijering/software/mtrackj/). We noted position of each retina frame by frame, beginning 8 seconds prior to the onset of sound as baseline, during the sound, and 8 seconds after the sound ended. For both left and right retinas, we measured the position of the central high acuity “elbow” and a point on the inside of the upper “arm” of each retina. Using these points, we analyzed two variables: how far the center point of each of the retinas moved and retinal angles, or torsion. We ran separate ANOVAs in JMP Pro (version 15.1) for movement and torsion. Time frame (before, during, or after sound playback), stimulus order, and their interaction were treated as fixed effects, with spider ID treated as a random effect. Tukey pairwise tests were used to distinguish between the three treatments.

We also tested whether the direction in which the principal eyes are directed is influenced by the position of the speaker that played the wasp sound. We used two independent variables in two different tests. First, because the speaker could be placed to the left, right, and behind the spider, and we wished to test whether the spider looked toward the speaker, we examined where the retinas moved along the horizontal axis. We subtracted the average *x* coordinates of the two retinas prior to sound playback from the average *x* coordinates of the two retinas during sound playback. A positive number indicated that the retina was, on average, further to the right during playback than prior to playback, and a negative number indicated that the retinas were, on average, further left during playback. Because we hypothesized that spiders might look upward when they hear a sound of an aerial predator, we also carried out a similar calculation for the vertical axis using the average *y* coordinates and tested whether spiders were more likely to look up when sound was played. We analyzed the data with a mixed-model ANOVA, with spider ID as a random effect, and stimulus order and position of the speaker as fixed effects.

### Experiment 2: Attentional shifts in response to predator sounds

For Experiment 2, we used a previously developed protocol to test whether sound influences spiders’ attention to a prey image. In this distraction protocol, we waited until a spider was actively scanning a primary image with its principal eyes with back-and-forth torsional movements of the retinas. We then positioned the distractor within the field of view of the forward-facing secondary eyes (the ALEs), outside of the field of view of the principal eyes as they examined the stimulus, but in an area of the screen that they could be redirected to see. Previous experiments showed that whether a spider looks at the distractor is influenced by both the nature of the primary image and of the distractor. In particular, spiders that are examining the initial image of a silhouette of a cricket are unlikely to direct their gaze to a distracting oval (Bruce et al., 2021). We used those same stimuli here. We predicted that spiders would be more likely to look at the oval if a wasp sound was played simultaneously with the oval’s appearance compared to trials in which no sound was played.

Trials began with a blank white screen. After a 5-min acclimation period, the initial stimulus appeared in the center of the screen. If the spider did not scan the stimulus within five minutes, the trial ended, and the spider was tested again at a later date. Spiders that examined the cricket did so by scanning their retinas over it, often accompanied by retinal vergence and torsion. All spiders were required to engage in at least two complete sequential back-and-forth scans before they were presented with the distractor. All individuals received both sound and no-sound treatments in random order (random.org/lists), and spiders were given a 3-minute rest period between treatments. In sound treatments, we presented the same sound and in the same manner as in Experiment 1, simultaneously with the appearance of the distractor. We tested 25 spiders.

Overlays were made using Adobe Premiere Pro 2019, which were aligned using an established calibration protocol (Bruce et al., 2021; Jakob et al., 2018). Each treatment was exported as a .mov file from Adobe Media Encoder and scored via video playback. We used a categorical scoring of responses: (1) the spider did not look away from the primary stimulus when the sound was played and the distractor appeared; (2) the spider shifted its gaze to the distractor oval and began scanning; (3) the spider shifted its gaze to the distractor but did not scan it; (4) the spider stopped scanning the primary stimulus but did not look at or scan the distractor, instead sweeping its retinas back and forth. For the primary analysis, we summed observations categories 2-4 into a category “looked away from initial stimulus.” In a second analysis, we used paired McNemar tests with Holm p-value corrections to test if any effects observed in the primary analysis were driven by the other component categories. Three trials could not be categorized due to video quality and were excluded from the analyses.

We analyzed the categorical retina movement binary response for each sound treatment using generalized linear mixed effect models (GLMMs) with binomial distributed errors using the ‘glmer’ function from the ‘lme4’ package (version 1.1-21, Bates et al., 2015) in R version 3.6.2 (R Core Team, 2020). Treatment type, sound or no sound, was treated as the fixed effect, while spider ID treated as a random intercept. Model fit and residual diagnostics were assessed with the ‘DHARMa’ package (Hartig and Hartig, 2017).

## RESULTS

### EXPERIMENT 1: RETINAL MOVEMENTS INCREASE IN RESPONSE TO PREDATOR SOUNDS

#### Do predator sounds impact eye movements?

In Experiment 1, *P. princeps* eye movements began immediately with sound onset, and continued after the sound ended. Unlike the methodical, torsional movement of scanning, retinas usually moved in rapid back-and-forth sweeps (Supplementary Video 2). Compared to when sound was absent, sound significantly increased the average distance the retinas moved from one frame to the next (*F*_2, 100_ = 85.20, *P* < 0.0001; Figure 2A). Stimulus playback order was not significant (*F*_1,100_ = 0.001, *P* = 0.9797). In Tukey’s follow-up tests, every pair significantly differed (*P* < 0.0001), with the least movement prior to the onset of playback and the greatest movement during sound playback. Similarly, retinas showed increased torsion with the onset of sound (*F*_2, 110_ = 58.09, *P* < 0.0001; Figure 2B). Tukey’s tests indicated that there were significant differences between each pair of treatments: retinal movements were lowest before the onset of the sound, increased during sound playback (before vs. during: *P* < 0.0001) and then decreased after sound playback (during vs. after: *P* < 0.05).

**Figure 2.**
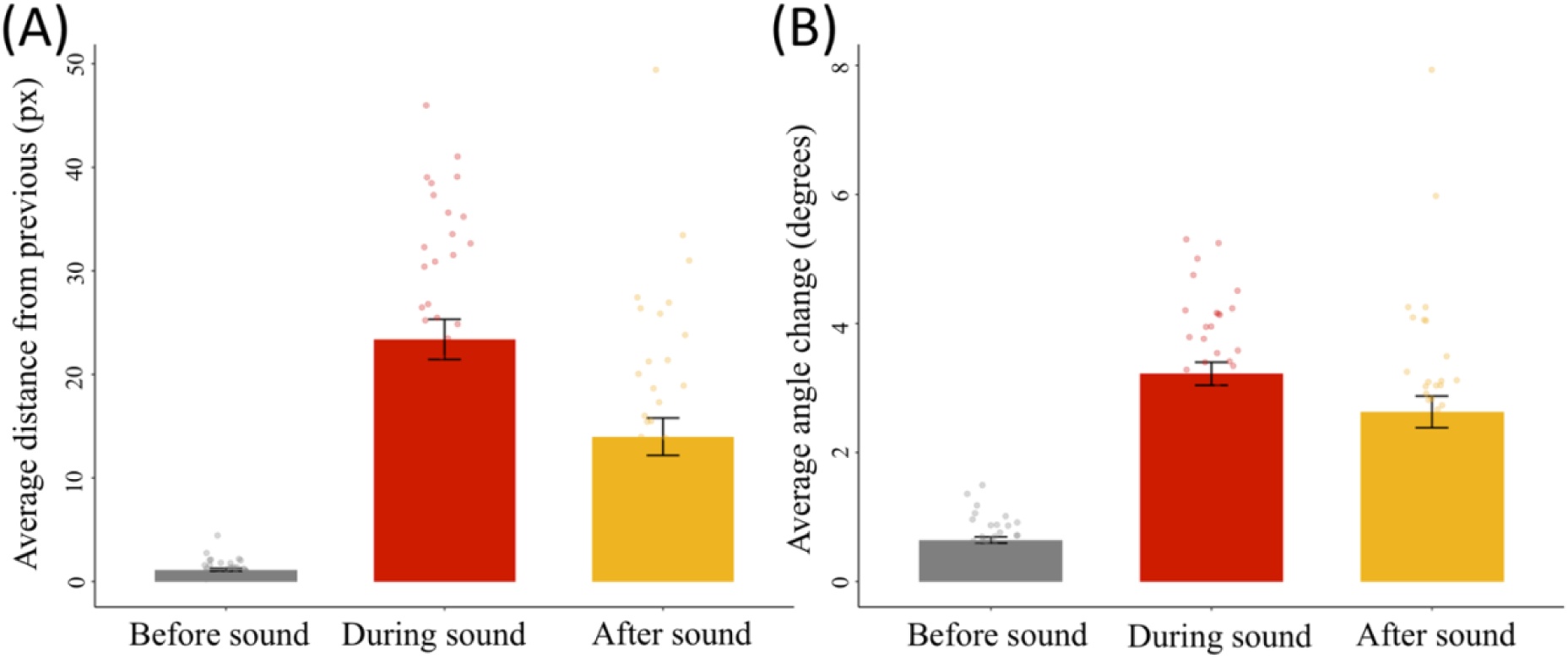
Predator sounds increase retinal activity. (A) The average distance traveled by the retinas increased during playback of predator sounds (red), which persisted even after playback had stopped (gold). (B) The average angle of retinal torsion also increased during and after predator sounds. The period of time measured before and after the stimulus was a total of 8 seconds each, while the buzz period lasted 5 seconds. In contrast to freezing their body posture, spiders move their eyes.

#### Do spiders look to the source of sound?

We found no effect of speaker position on retinal position. The horizontal position of the retina was not influenced by speaker placement (*F*_2,23_ = 0.233, *P* = 0.79) or stimulus order (*F*_1,23_ = 1.22, P = 0.28). Similarly, the vertical position of the retina was not influenced by speaker placement (*F*_2,23_ = 1.139, *P* = 0.34) or stimulus order (*F*_1,23_ = 1.370, *P* = 0.25). Taken together, these results suggest that sound disrupted spiders’ resting position, as evidenced by increased retinal movements, but we found no evidence that spiders were orienting to the source of the sound.

### EXPERIMENT 2: ATTENTION IS DISRUPTED IN RESPONSE TO PREDATOR SOUNDS

#### Do predator sounds cause spiders to shift their gaze away from the initial stimulus?

Spiders shifted their gaze away from the initial salient stimulus (responses 2-4 combined) significantly more frequently with wasp sound than when sound was not present (Figure 3; Supplementary Table 1; GLMM: 2.0531 ± 0.9148, *z* = 2.244, *P* = 0.025).

**Figure 3.**
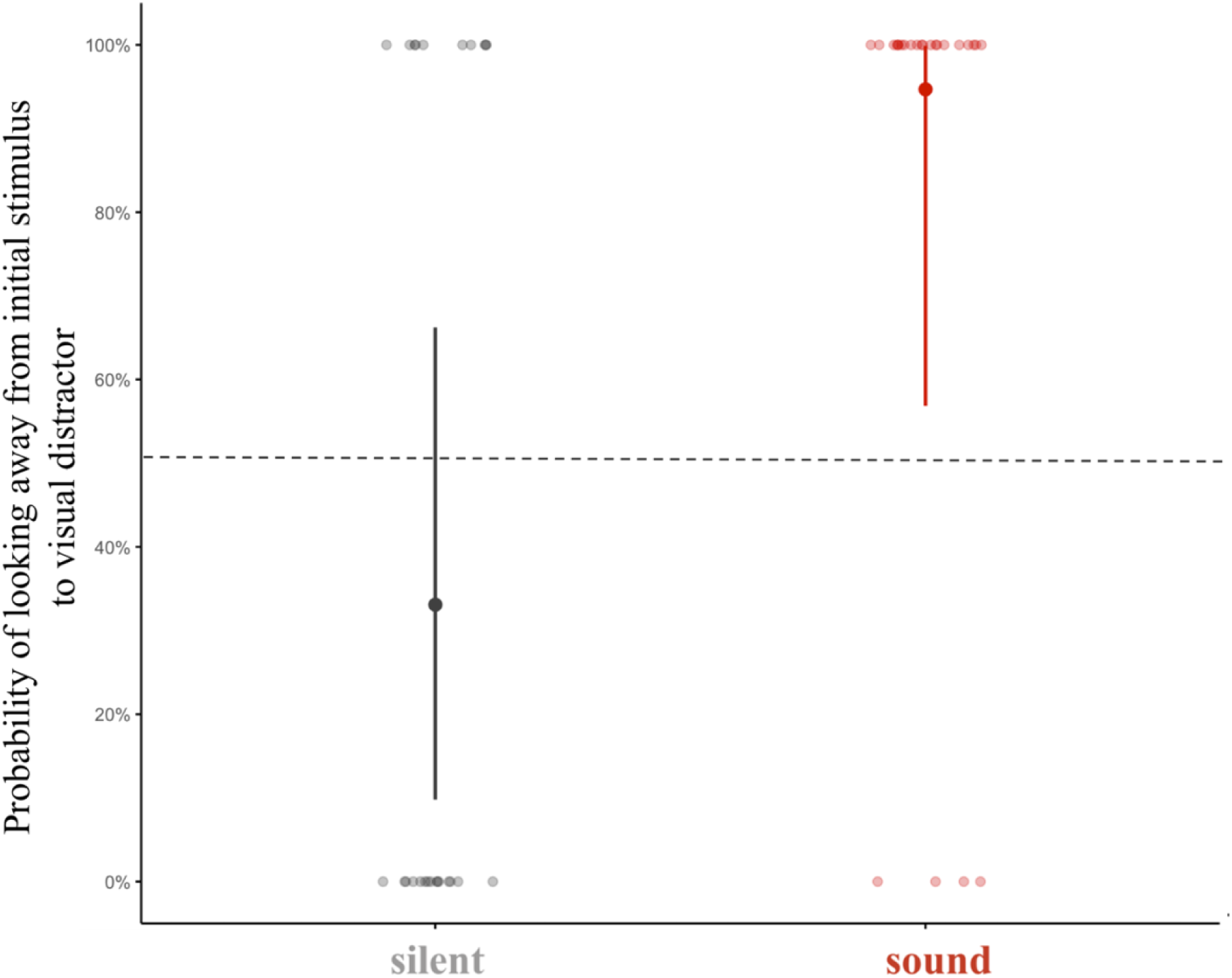
Predator sounds disrupt attention. With sound exposure, spiders were much more likely to look away from the initial stimulus (right) compared to silent controls (left). Bars show GLMM-estimated probabilities with their 95% confidence intervals. The dotted line represents chance, and the faint points represent binary choices of each individual spider.

#### Do spiders look to an appearing visual stimulus?

Spiders were not more likely to look at the distractor when sound was played (Supplementary Figure 1; Supplementary Video 3). Instead, spiders distracted from the initial stimulus by sound moved their eyes back and forth in an erratic sweeping motion rather than directing their gaze to the distractor stimulus (Supplementary Table 2; *McNemar* tests with *Holm* correction: “did not look away” reactions decreased with sound exposure, P = 0.008, whereas “undirected movement” increased with sound, P = 0.023; redirecting to the oval distractor and oval scanning did not differ between conditions P ≥ 0.4). Similar to Experiment 1, these results show that spiders start sweeping eye movements in response to wasp sound, but in doing so disengage with a visual stimulus and simultaneously ignore a distractor.

## DISCUSSION

As predicted, spiders moved their principal eyes more in response to sound. At the onset of the sound of a predatory wasp, spiders immediately swept their retinas from side to side, also increasing their torsional retinal movements (Experiment 1). Shamble et al., (2016) previously showed that spiders freeze when they detect airborne wasp sounds. Taken together with our finding of increased retinal movement, this suggests that spiders are able to decouple eye movements from body movements when responding to potential threats. Predatory wasps are documented to be voracious predators of salticids (Edwards, 2004), and mud daubers of the genus *Trypoxylon* hover close to branches, leaves, and other surfaces while hunting in flight, where they are fast and agile in the air but comparatively slow and awkward on the ground (Coville, 1987). Shamble et al. (2016) further noted that broadband sounds similar to wasp flight may also signal other dangers, such as predators approaching through leaf litter. In this context, remaining still while actively moving the eyes may allow spiders to visually sample their surroundings for potential danger without revealing their position. We therefore reasoned that increased retinal movement could provide a means for spiders to search for threats while minimizing the risk of detection.

Despite the clear increase in eye movements, we found no evidence that the direction of eye movements was influenced by the spatial location of the speaker. One possibility is that spiders without a web have difficulty localizing airborne sound, which could lead them to sweep their retinas without successfully identifying a source. Another possibility is that the sound caused the ball to vibrate, which could have obscured directional cues. Jumping spiders likely detect airborne sounds using airflow sensitive hairs called trichobothria on their legs (Shamble et al., 2016). The tuning and spacing between these hairs are unlikely to be sufficient for comparing inputs in order to determine sound direction. However, another mechanosensory structure, the lyriform organ, is strain sensitive but also responsive to sound, and lyriform organs are distributed across all eight legs (Stafstrom et al., 2020), raising the possibility that it could contribute to sound localization. Given our results, it may be more plausible that retinal sweeping does not function to localize the sound source at all. Instead, it may reflect a generalized startle response, an attempt to identify potential escape routes, or a consequence of the fact that large, nearby sounds alone do not provide reliable spatial information about the location of a threat.

While spiders did increase their principal eye movements in response to sound, we did not find that they were better at directing their gaze to the location of a suddenly appearing visual stimulus (Experiment 2). When spiders were examining an image of a cricket, the onset of a wasp sound made them more likely to look away from the cricket, indicating that sound disrupted ongoing visual attention, much like how sound also disrupted spiders at rest. We had predicted that spiders inspecting the cricket would be more likely to direct their gaze toward a distractor stimulus that appeared at sound onset, as they might interpret the new visual stimulus as the source of the sound. Instead, spiders typically engaged in the same erratic retinal sweeping behavior described previously rather than orienting toward the new stimulus. This pattern is consistent with findings in primates, where abrupt sounds can disrupt ongoing visual orienting. For example, in humans, unexpected sounds can briefly interrupt reading by disrupting eye movements across the text and drawing attention away from it (Vasilev et al., 2019). Such responses may simply be a startle reflex, or instead reflect an adaptive strategy under conditions of high uncertainty, in which committing to any single visual target could delay escape or interfere with responding to a subsequent, higher priority stimulus.

The choice of stimuli may also have influenced these results. We selected an oval because we wanted a distractor that by itself was not particularly attractive to the spider so that we could more easily detect an effect of the sound. For example, in previous experiments, an appearing oval was more likely to attract the gaze of spiders that were already examining a different oval compared to those that were examining a more salient cricket, while a looming stimulus always attracted the gaze of the principal eyes (Bruce et al., 2021). It is possible that spiders in the current experiment may have shifted their gaze to a different distractor more readily, or perhaps may have looked more to a congruent stimulus such as a wasp silhouette. Spiders might also have responded differently to the wasp sound if it were acoustically looming (Ghazanfar et al., 2002). The timescale of jumping spiders’ retinal sweeping movements in response to wasp sounds is longer than is typically associated with startle reflexes alone yet appears insufficient to trigger prolonged surveillance behavior. Future work will be needed to fully disentangle to what extent wasp sounds induce retinal sweeping as a bottom-up reflexive startle response, or a temporary top-down state of heightened vigilance, as well as whether such exposure subsequently prompts other defensive behaviors, such as seeking shelter.

Overall, our results demonstrate that cross modal cues can disrupt attention in jumping spiders. When considered alongside our previous findings of visual distraction (Bruce et al., 2021) and arousal (Winsor et al., *unpublished*), these results suggest that jumping spiders exhibit a rich repertoire of attentional responses across sensory modalities. Remarkably, these responses parallel those observed in much larger-brained vertebrates that evolved independently, pointing to broad evolutionary convergence in oculomotor constraints and attention.

## Acknowledgments

We thank Ethan Clotfelter, Sarah Partan, Lisa Mangiamele, Laela Sayigh, Renae Brodie, Jeff Podos, Luke Remage-Healy, and Paul Katz for providing comments on drafts of the manuscript. This research was funded by NSF IOS 1656714 to E.M.J. and R.R.H.

## SUPPLEMENTARY INFORMATION

**Supplementary Figure 1.**
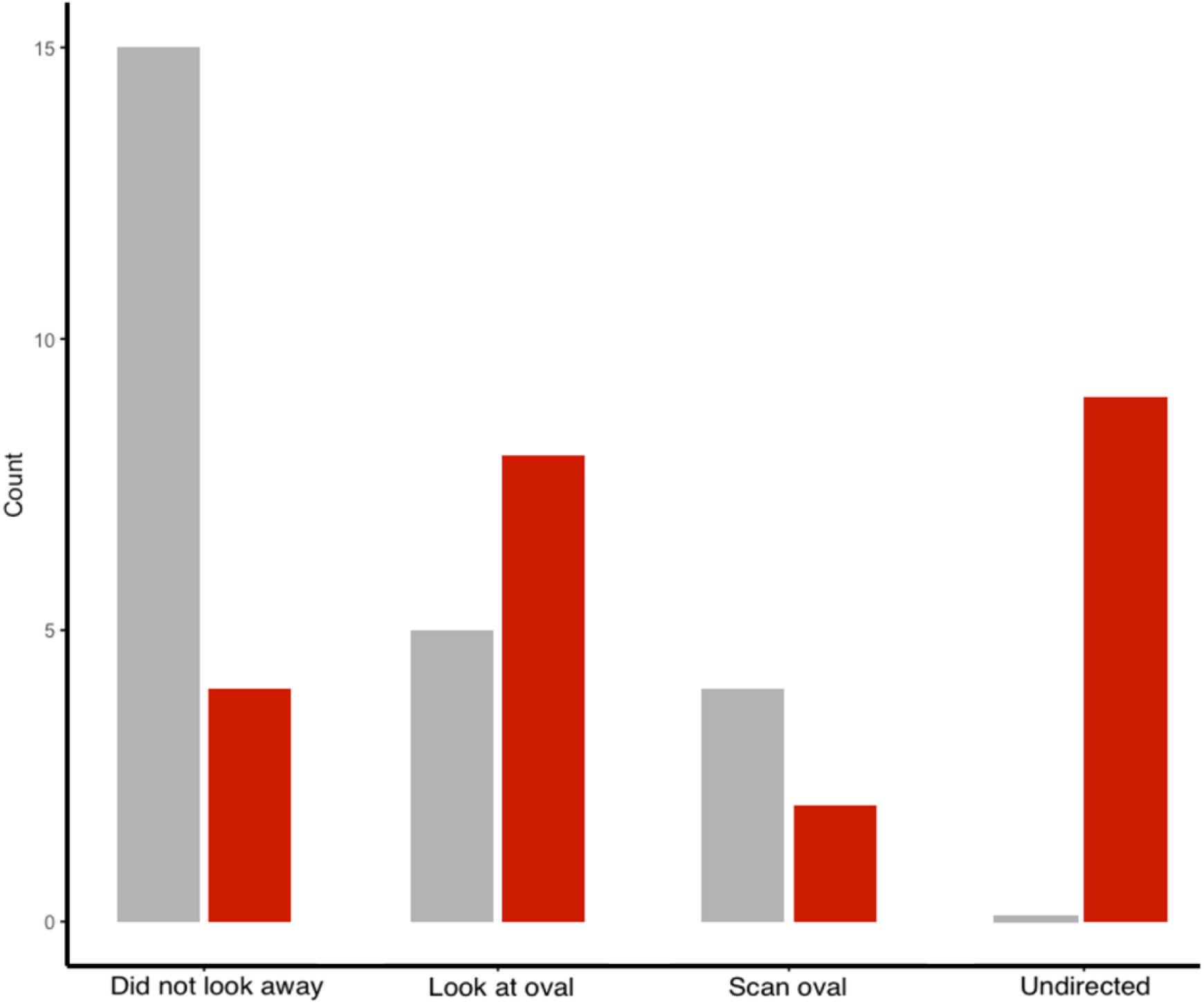
Sound-exposed spiders were not more likely to look at the oval distractor after disengaging with the initial cricket stimulus. Control spiders (grey) were more likely than sound-exposed spiders (red) to not look away and hold their attention on the initial cricket stimulus. However, we did not see strong evidence that sound-exposed spiders were more likely to look at, or especially scan, the distractor oval; we instead found that sound-exposed spiders exhibited greater undirected movement (i.e., sweeping retinal movements that were disengaged with all visual stimuli, a pattern we did not observe in control spiders). Three trials were excluded due to video quality. Bars show raw counts of responses across all spiders.

**Supplementary Table 1.** GLMM model estimates (±SE) of the probability of a spider disengaging with a cricket stimulus after sound exposure. Values are relative to the intercept which represents a control with no sound. Z and P values are indicated for each fixed effect. The bolded significant P value shows that sound-exposed spiders were significant more likely to look away from the initial cricket stimulus than were control spiders.

| Factor | Estimate | ±SE | z | P |
| --- | --- | --- | --- | --- |
| Intercept (Control) | -0.437 | 0.437 | - | - |
| Treatment is sound | 2.053 | 0.915 | 2.244 | **0.025 \*** |

**Supplementary Table 2.** Changes in response types with sound exposure. Paired McNemar tests examining changes in spiders’ initial reactions between no-sound and sound conditions. Values indicate the number of individuals (N = 22) that switched into (“gained”) or out of (“lost”) each reaction category when sound was present. Three spiders that exhibited responses in a trial that could not be scored within these categories were dropped from the analysis. Significant P values, Holm-corrected for multiple comparisons, are bolded. Sound exposure significantly decreased “did not look away” reactions and significantly increased “undirected movement” reactions, whereas redirecting to the oval distractor and oval scanning reactions did not differ between conditions.

| Reaction type | N (paired by individual ID) | “Gained” with sound exposure | “Lost” with sound exposure | P (Holm-corrected) |
| --- | --- | --- | --- | --- |
| Did not look away | 22 | 0 | 10 | **0.008 \*** |
| Undirected movement | 22 | 8 | 0 | **0.023 \*** |
| Redirect only | 22 | 5 | 1 | 0.438 |
| Oval scanning | 22 | 2 | 4 | 0.688 |

## SUPPLEMENTARY VIDEOS

https://drive.google.com/file/d/1d2M7JqUXs6JAomq-fWAIgC3JLf_fIf55/view?usp=drive_link

**Supplementary Video 1. The buzz sound played to spiders across both experiments.** The buzz sound used a broadband range of frequencies. Spectrogram video was created with the open-source Spectrum Analyzer software (Academo, 2025).

https://drive.google.com/file/d/1HYWgCzGTo9THq5NKvbxT1s_dpp3oIvxO/view?usp=drive_link

**Supplementary Video 2. Eye movements in response to predator sounds.** The retinas sweep in a stereotypical rapid, back-and-forth sweeping motion.

https://drive.google.com/file/d/1mVtgvkW9x-5kiLe6h1U53E0yWsyz5Uu5/view?usp=drive_link

**Supplementary Video 3. Spiders disengage with all stimuli in response to predator sounds.** The retinas disengage with the initial cricket stimulus, but instead of shift toward the oval distractor stimulus, the retinas move in the same stereotypical rapid, back-and-forth sweeping motion.

## Notes

### Competing Interest Statement

The authors have declared no competing interest.

